# Virus-host interactome reveals host cellular pathways perturbed by tick-borne encephalitis virus infection

**DOI:** 10.1101/2025.02.26.640292

**Authors:** Liyan Sui, Ying-Hua Zhao, Heming Wang, Hongmiao Chi, Hanxi Xie, Tian Tian, Wenfang Wang, Mucuo Zhu, Naicui Zhai, Zhixia Song, Yueshan Xu, Kaiyu Zhang, Lihe Che, Guoqing Wang, Nan Liu, Yicheng Zhao, Quan Liu

## Abstract

Tick-borne encephalitis virus (TBEV) poses an increasingly significant threat to public health. Here, we delved into the virus-host interactome and discovered that TBEV has a profound impact on multiple host cellular pathways. Viral pre-membrane protein (prM), non-structural proteins NS1, NS2B3, NS4A, and NS5 remodel the host’s innate immune responses. PrM and NS2B3 are involved in autophagy, NS4A is associated with neurodegenerative diseases, and NS5 participates in spliceosome dynamics, ribosomal biogenesis, cell cycle regulation, and DNA damage response. Notably, TBEV infection causes G0/G1 cell cycle arrest in host cells. NS5 interacts with histone acetyltransferase P300 to upregulate P16 expression, suppressing CDK4 and resulting in cell cycle arrest. Elevated P16 and reduced CDK4 levels were observed in TBEV-infected brain organoids. The P300 inhibitors and CDK4 agonist can reverse virus-induced cell cycle arrest and inhibit viral replication. Further analysis uncovered potential antiviral targets like KAT6A, XIAP, and RIOK1/BRD9. These findings provide valuable insights into TBEV pathogenicity and hold promise for antiviral drug development.

## Introduction

Tick-borne encephalitis virus (TBEV), a zoonotic flavivirus within the *Flaviviridae* family, is transmitted by ticks and ranks among medically significant pathogens such as dengue fever virus (DENV), Zika virus (ZIKV), yellow fever virus (YFV), West Nile virus (WNV), and Japanese encephalitis virus (JEV) (*1–3*). Endemic to Europe and northeastern Asia, TBEV causes severe neuroinvasive infections that may progress to meningitis or encephalitis by targeting the central nervous system (CNS). Although inactivated vaccines are clinical available, no specific antiviral therapies exist for TBEV infections. Emerging challenges, including genetic mutations and vaccine breakthrough cases (*4–6*), highlight the critical need to elucidate the pathogenesis of TBEV and develop novel antiviral strategies.

The TBEV genome comprises a single-stranded, positive-sense RNA of approximately11 kb, encoding three structural proteins: capsid (C), envelope (E) and pre-membrane (prM) proteins, which mediate viral entry, assembly and package (*7, 8*). Additionally, seven non-structural proteins (NS1, NS2A, NS2B, NS4, NS4A, NS4B, and NS5) orchestrate viral replication by interacting with host factors (*9, 10*). For instance, NS2B forms a protease complex with NS3, while NS5 disrupts type I interferon signaling by binding host proteins Tyk2 and Scribble, thereby enhancing viral replication (*11, 12*). Although studies on individual viral protein have advanced our understanding of TBEV pathophysiology (*13–17*), the molecular mechanisms governing host cell responses to infection remain poorly characterized, hindering insight into viral pathogenicity and the identification of therapeutic targets. Investigating virus-host interactions can reveal complex networks of host proteins subverted by viral proteins (*18, 19*). In this study, we mapped the virus-host interactome and uncovered its profound impact on critical host cellular pathways, including innate immunity, autophagy, ribosomal biogenesis, cell cycle regulation, and DNA damage response. These findings elucidate molecular strategies underlying TBEV pathogenicity and provide a foundation for developing targeted antiviral therapies.

## Results

### Analysis of TBEV-host protein interactions

Identifying virus-host interactions can be valuable for understanding how viruses hijack and re-wire host machinery. We conducted an extensive analysis of protein-protein interactions between TBEV and host. TBEV proteins, each tagged with a 2xStrep affinity, were introduced into HEK293T cells. All ten viral proteins, along with the additionally constructed NS2B3 (comprising NS2B and NS3 to mimic an active protease) were correctly expressed and subjected to affinity purification and mass spectrometry (AP-MS) analysis (Figures 1a-c, S1; Table S1).

**Figure 1.**
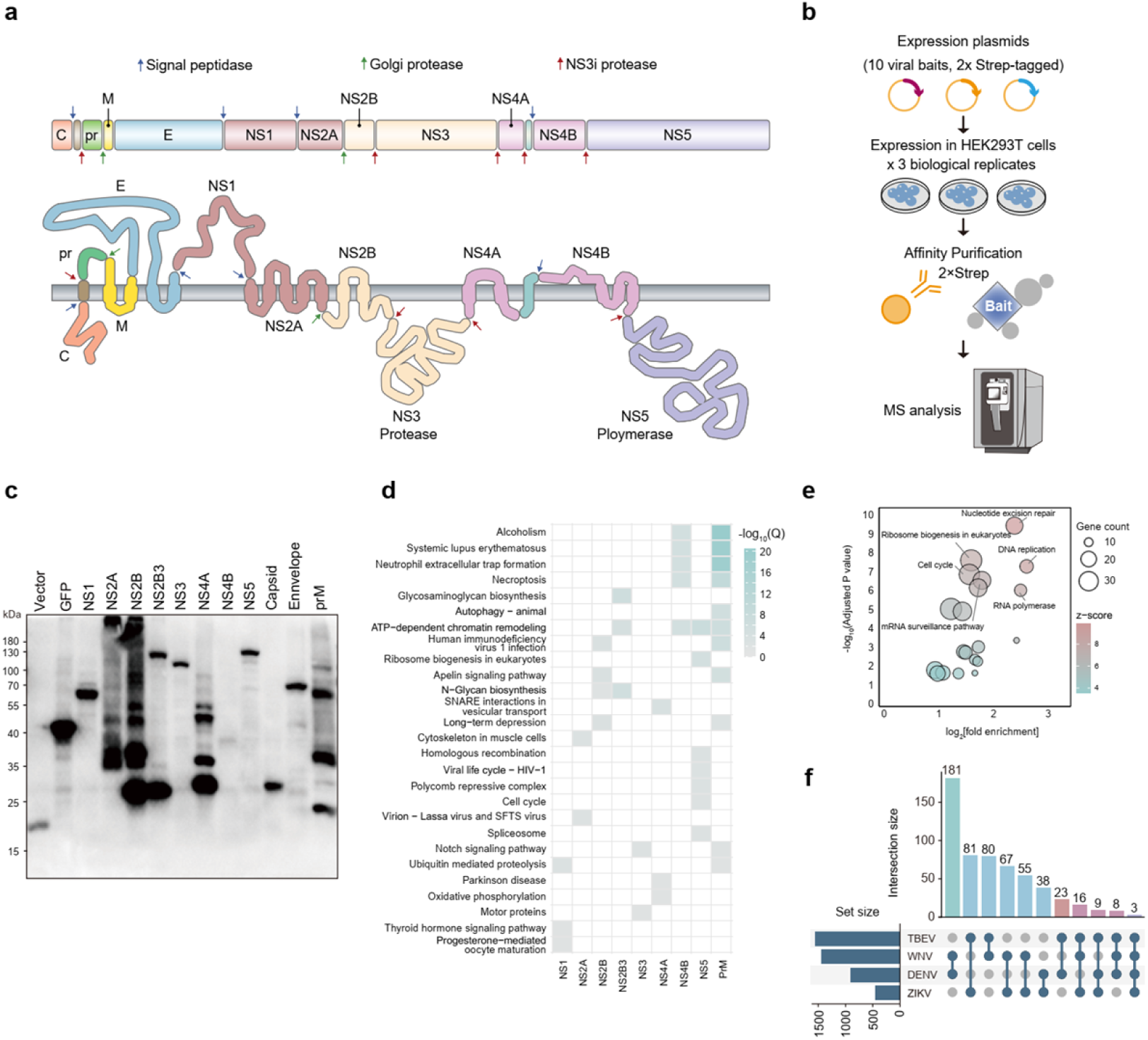
Summary of TBEV-host protein-protein interaction network. **a.** Schematic view of TBEV viral proteins. **b.** Comprehensive interactome analysis was conducted to study interactions between TBEV and host proteins. **c.** Viral baits expressed in HEK293T cells and analyzed by immunoblot assay using anti-Flag antibody. **d, e.** Enrichment analysis of TBEV-interacted host proteins (d) and the proteins (e) significantly deregulated by TBEV infection. **f.** UpSet plot showing the overlap of flavivirus-host protein–protein interactions from other studies.(*18, 19*) Each bar shows the interactions shared by only the marked studies at the bottom.

Virus-host interaction network analysis identified 2201 TBEV-host protein-protein interactions and revealed a wide range of cellular activities intercepted by TBEV (Figures 1d; Table S1). These encompassed inflammatory processes including systemic lupus erythematosus, neutrophil extracellular trap formation, and the infection of other viruses, such as HIV-1, Lassa virus and severe fever with thrombocytopenia syndrome bunyavirus (SFTSV). The cellular processes that play important roles during virus infection were also enriched in TBEV-interacted proteins, such as autophagy and cell cycle, as well as cellular metabolism, with a focus on glycosaminoglycan biosynthesis and N-glycan biosynthesis. Additionally, TBEV-host interactions extended to necroptosis and ribosome biogenesis (Figures 1d). Consistently, our analysis of proteins significantly affected by TBEV infection also uncovered an enrichment in pathways associated with ribosome biogenesis, cell cycle, and mRNA processing, suggesting that these pathways might be regulated through interactions with the virus (Figures 1e; Table S2) (*20*).

Furthermore, we conducted a comparative analysis of our TBEV interaction map with other flaviviruses, including DENV, ZIKV, and WNV, and found that 81 preys were common between TBEV and ZIKV, while 23 and 80 preys were shared between TBEV with DENV and WNV, respectively (Figure 1f; Table S3). Moreover, there were 3 preys (HDGFL2, GOLT1B and G6PC3) present in all four flaviviruses, underscoring the existence of conservative regulatory mechanisms among various flaviviruses.

Notable, viral proteins-interacted host proteins were significantly enriched with biological processes consistent with their known biology and subcellular location (Figure 2a). For example, NS5 was enriched for interacting with proteins linked to nuclear functions, including cell cycle, chromatin remodeling and ribosome biogenesis, aligning with its nuclear location (Figure S2). NS2B and NS2B3 targeted components are associated with ER-associated degradation (ERAD), such as NS2B-SYVN1, NS2B-HERPUD1 and NS2B3-MAN1B1. This interaction may be associated with the aggregation of ER in cells overexpressing NS2B and NS2B3 (Figure S2). We also observed the enriched autophagic process of prM interacting proteins, such as AKT1, VPS11 and VPS41. NS4A was notably associated with proteins implicated in neurodegenerative diseases, including Parkinson’s, with enrichment of proteins such as ND3 and SLC39A7. This finding aligns with the role of NS4A in ZIKV, highlighting a conserved function of NS4A in the neuropathogenesis of flaviviruses (*21*). prM and NS1 were also found to interact with proteins involved in ubiquitin-mediated proteolysis, such as UBE3A and CUL7, which play important roles during virus infection (*22, 23*).

**Figure 2.**
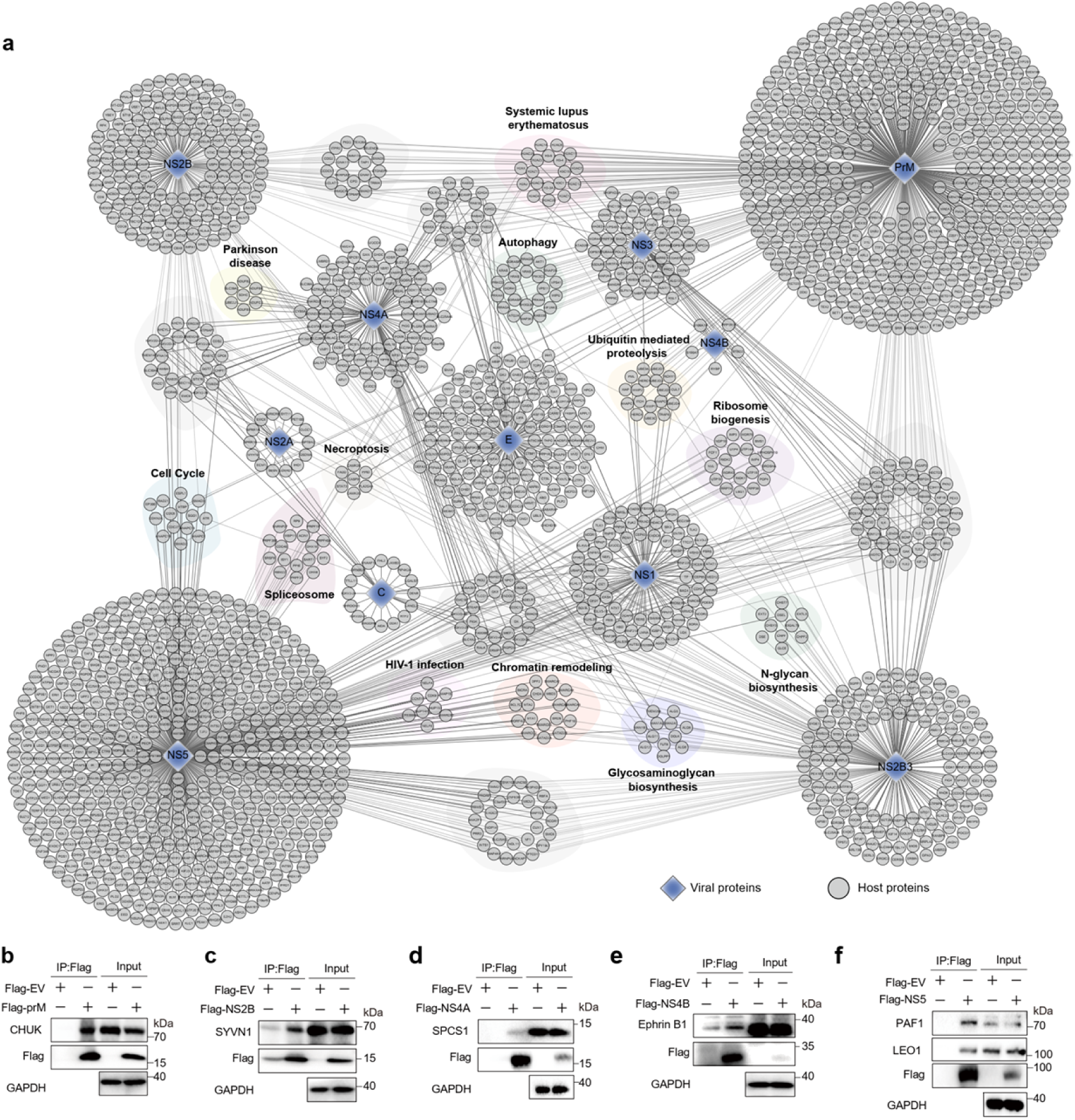
The TBEV-host interactome. **a.** An integrated protein-protein interaction network showed high-confidence TBEV-host interactions. Viral proteins are represented as rhombus nodes. Edge intensity reflects the MIST value associated with the interaction. Solid thick edges represent the most significant interactions, while light edges correspond to low MIST (Molecular Interaction Search Tool) value (0.5 < MIST < 0.99). **b-f.** Validation of virus-host interactions using co-immunoprecipitation. Flag-tagged empty vector (EV) or viral proteins were overexpressed in HEK293T cells, and CHUK co-precipitated with prM (**b**), SYVN1 co-precipitated with NS2B (**c**), SPCS1 co-precipitated with NS4A (**d**), endogenous Ephrin B1(EFNB1) co-precipitated with NS4B (**e**), along with PAF1 and LEO1 co-precipitated with NS5(**f**), were detected using flag beads and immunoblot assay.

In particular, we identified previously reported proteins that showed bona fide interaction with proteins from other flaviviruses, including DENV and ZIKV NS5 with PAF1C complex (PAF1 and LEO1) (*19, 24*), JEV NS2B with signal peptidase complex subunit 1 (SPCS1) (*25, 26*), and ZIKV prM with mitochondrial antiviral signaling protein (MAVS) (*15*), confirming these processes as important targets of diverse flaviviruses (Figures 2a). The interactions of prM- CHUK, NS2B-SYVN1, NS4A-SPCS1, NS4B-EFNB1, NS5-PAF1 and NS5-LEO1 were further validated by co-immunoprecipitation analysis (Figures 2b-f), demonstrating the credibility of our analysis.

Building on our prior work demonstrating that TBEV infection significantly alters host protein phosphorylation and acetylation (*20*), we expanded this analysis by mapping perturbed host proteins onto TBEV the interactome. A substantial subset of viral proteins exhibited associations with host proteins, suggesting that viral proteins may directly influence the modifications of host proteins (Figures S3 and S4). For instance, while DENV and ZIKV NS5 proteins are known to interact with LEO1 and PAF1 to suppress interferon-stimulated genes (ISGs) expression and evade immunity (*19*), our study revealed that TBEV infection upregulated the phosphorylation of LEO1 at S607 and S608, sites critical for its transcriptional activity (*27*). This finding implies a distinct yet functionally convergent mechanism by which flaviviruses may manipulate host machinery to subvert antiviral responses.

### Key cellular pathways perturbed by TBEV infection

To further understand the signaling pathways disrupted by TBEV, we applied an automated approach to explore the underlying molecular mechanisms within the viral interactome and proteome(*20*). We integrated viral-host interactions and perturbed protein abundance into the global human interaction network and utilized a network diffusion approach. This analysis employed known cellular protein–protein interactions, signaling and regulation events to identify connection points between cellular proteins interacting with viral proteins and those affected by TBEV infection (Figure 3a, S5). NS5 was shown to participate in spliceosome, DNA damage response (DDR), and cell cycle progression. Meanwhile, prM can modulate autophagy. These functions, along with the regulation of the innate immune response and ribosomal biogenesis by other viral proteins, highlighted the multifaceted roles of TBEV proteins in host cell biology (Figures 3b-3e, S6 and S7).

**Figure 3.**
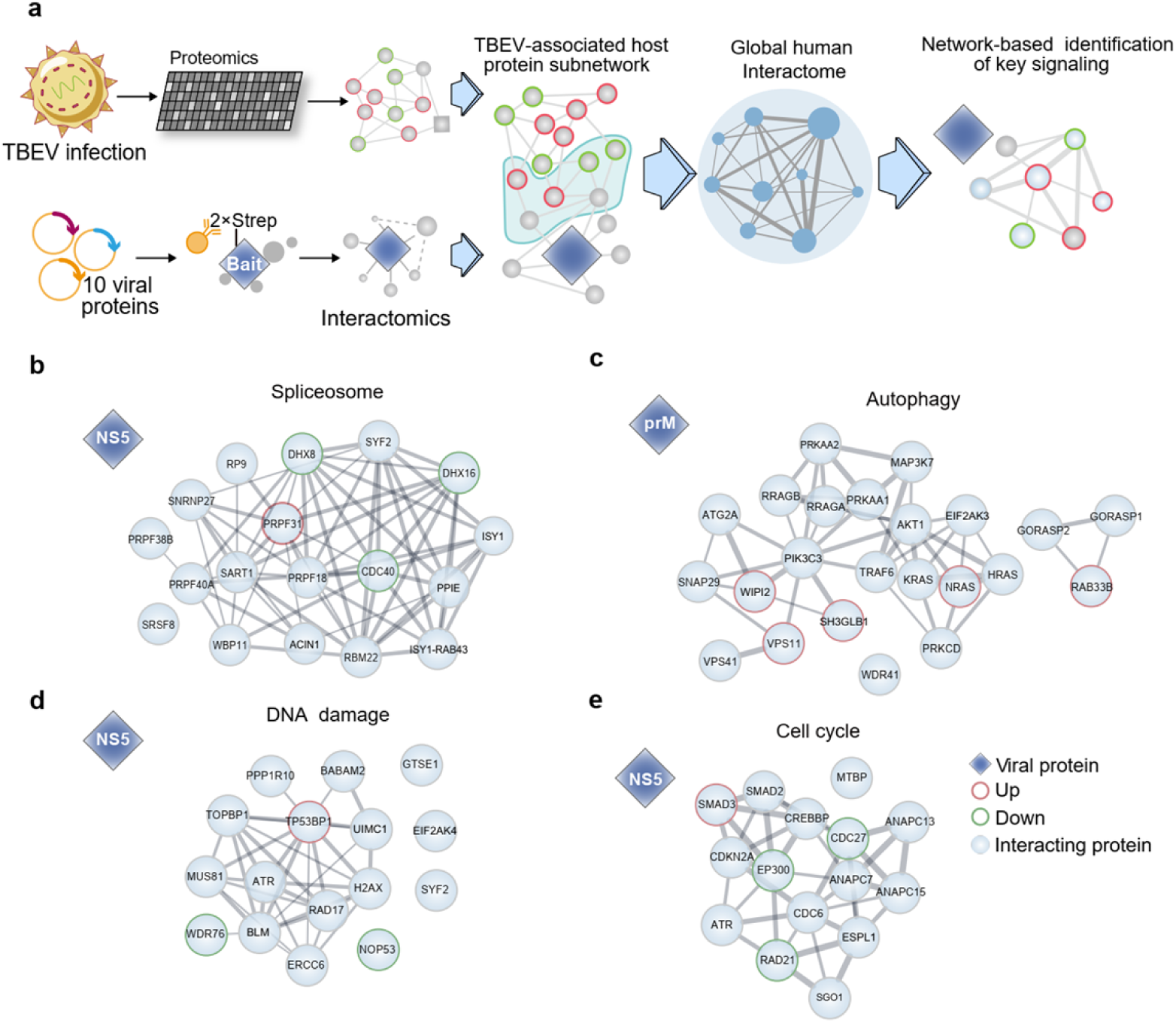
The key cellular pathways perturbed by TBEV. **a.** The schematic illustration outlines the network diffusion analysis process. The proteome results were initially integrated with the virus-host interactome, and subsequently mapped to the global human protein-protein interactome to identify affected host pathways. **b.** Subnetworks derived from network diffusion predict the correlation of TBEV NS5 with spliceosome (| Log_2_FC|>0.1, P value < 0.05). **c.** Subnetworks resulting from network diffusion depict the correlation of TBEV prM with factors involved in autophagy (|Log_2_FC|>0.1, P value < 0.05). **d, e.** Additional subnetworks derived from network diffusion predict the correlation of TBEV NS5 with DNA damage response (**d**) and cell cycle (**e**) (|Log_2_FC|>0.1, P value < 0.05). Black edges denote connections present in ReactomeFI. The thickness of the line reflects the combined score provided by STRING. The thickness of directed edges is proportional to the random-walk transition probability.

TBEV infection activates host cell interferon signaling, primary through upregulation of TLR3 and DDX58 (RIG-I). The elevation of RIG-I levels correlates with TRIM25 acetylation at lysine 345, a modification induced by TBEV infection. Furthermore, TLR2 activation, together with IKKα (CHUK) downregulation mediated by prM-CHUK interaction, leads to the increased expression of NFKB1 and NFKB2 (Figure S6a). To counteract host defenses, TBEV encodes multiple viral proteins to antagonize innate immune pathways. prM and NS1 interact with IRAK1, while NS4A binds ESCIT, collectively inhibiting NK-κB signaling. Simultaneously, prM targets TRIM4 and MAVS, while NS2B3 associates with UNC93B1, effectively blocking interferon production. NS5 directly binds to Tyk2 and SETD2, impairing interferon signaling and antiviral responses (Figure S6a).

TBEV infection has been shown to reduce synthesis of ribosomal RNA (rRNA) and its precursor (*28*). Our network analysis revealed a connection between NS5 and H/ACA ribonucleoprotein complex subunit 3 (Nop10), as well as the 90S pre-ribosome components (UTP14, UTP15, Rix7, Imp3 and Imp4). Intriguingly, we noted a decrease in the expression of Rrp7 and EMG1, accompanied by increased phosphorylation events on UTP3 (S464), UTP6 (S286), UTP8 (S212), and Rix (S515) (Figure S6b). Simultaneously, TBEV NS5 was found to interact with spliceosome-related proteins, such as CACTIN, DHX8, DHX16 and CDC40, which may be responsible for the downregulation of their expression and the remodeling of host RNA splicing (Figure 3b).

Our study indicated that prM, NS1, NS2B3 and NS4B could participate in the modulation of autophagy induction (Figure 3c, S7a). Consistent with our previous findings (*20*). prM was shown to activate autophagy via interacting with AKT1, NS2B3 and prM regulated autolysosome formation through binding to STX17 and VPS11, respectively. Additionally, NS1 and NS4B were also shown to regulate the activation of autophagy via interacting with S2K3 and ATG7.

TBEV infection was also implicated in triggering DDR, primarily induced by NS2B3 and NS5 through their interactions with DNA damage sensors, such as RAD17 and ATR (Figure 3d, S7b). Additionally, we noted a reduction in the expression of RRM2, a phenomenon previously observed during SARS-CoV-2 infection, which has been associated with DNA damage induction (*29*). Moreover, NS5 was implicated in the regulation of DNA damage repair through its interaction with SIRT1 to regulate the acetylation of Ku70 and KAP1 (Figure 3d, S7b). One consequence of DNA damage is cell cycle arrest. Our analysis suggested that NS5 may interact and regulate the expression of factors associated with mitosis, G1/S, and G2/M transition (Figure 3e, S7c), implying a role in the modulation of cell cycle progression during TBEV infection.

### TBEV infection induces G0/G1 cell cycle arrest to promote viral replication

Given the strong indications of cell cycle involvement during TBEV infection, we set out to investigate the impact of TBEV infection on cell cycle progression. Vero cells infected with TBEV at varying multiplicities of infection (MOI) of 0.1, 1.0 and 10 were analyzed for cell cycle dynamics. In the mock-infected group, 61.12% of cells were in the G0/G1 phase, a percentage that significantly increased to 67.48%, 71.03%, and 71.04% in MOI 0.1, 1.0 and 10 TBEV-infected groups, respectively (Figure 4a, b). To validate these findings, we extended our analysis to A549 and T98G cells that are susceptible to TBEV infection (*30*), showing obvious elevation of G0/G1 cells upon TBEV infection at an MOI 10 (Figure S8a-d), underscoring the ability of TBEV infection to induce G0/G1 cell cycle arrest.

**Figure 4.**
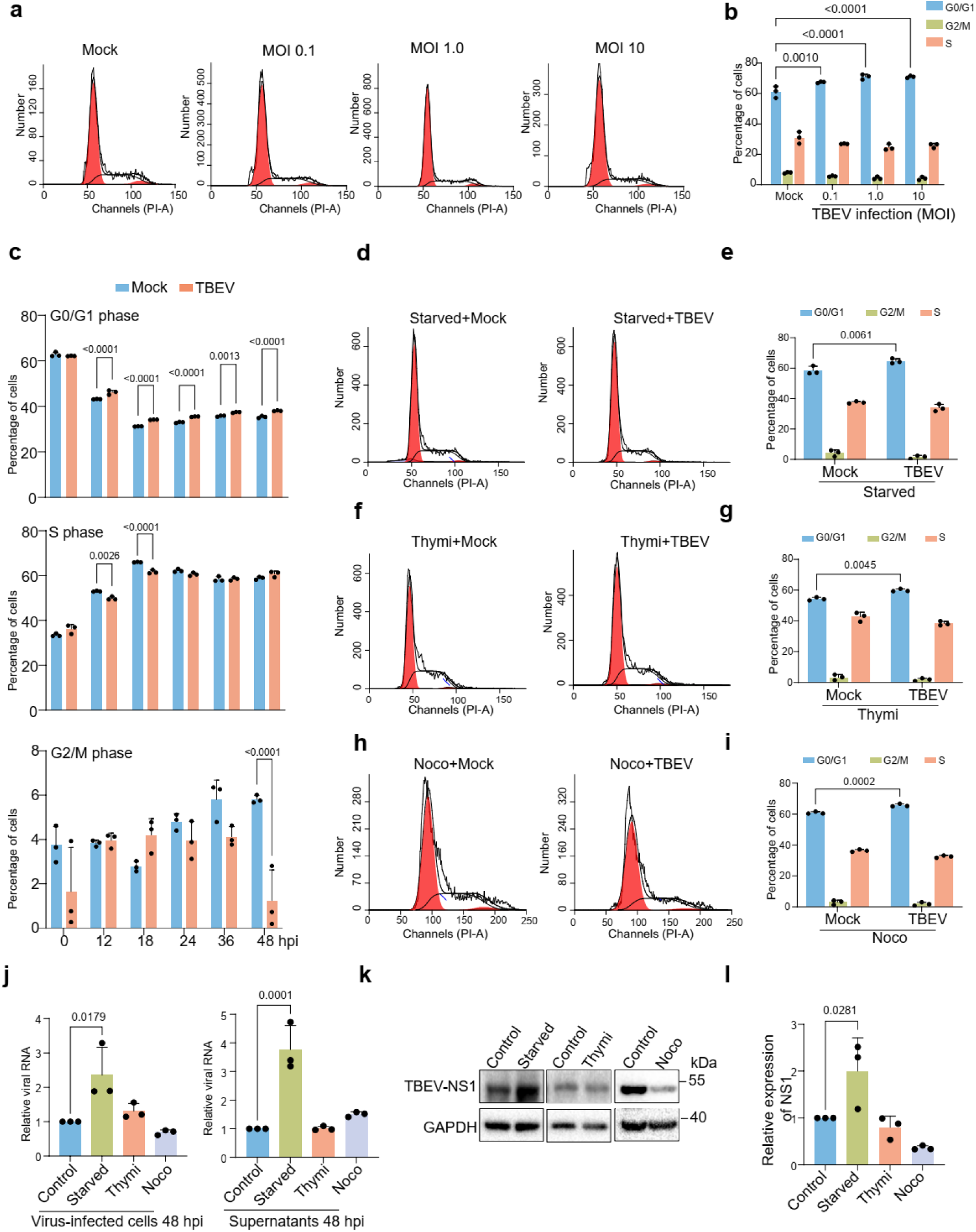
TBEV infection induce G0/G1 cell cycle arrest to promote viral replication. **a.** Vero cells were either mock-infected or infected with TBEV at various MOI (0.1, 1.0 and 10), cells were collected at 48 hours post infection (hpi) and their distribution was analyzed by flow cytometry. **b.** The experiments in (**a**) were repeated for three times, and the proportions of cells at different cell cycle phases were shown in column graph. **c.** Vero cells were mock-infected or infected with TBEV at the MOI of 1.0, cells were collected at 12, 18, 24, 36 and 48 hpi, and the percentages of cells in G0/G1 (*up*), S (*middle*) and G2/M (*down*) phase were analyzed by flow cytometry. **d-i**. Vero cells were synchronized to G0/G1 (**d**), S (**f**) and G2/M (**h**) phases via no-serum starvation, adding 0.85mM Thymi or 25 ng/mL Noco for 20 h. Subsequently, the cells were either mock-infected or infected with TBEV at an MOI of 10, cells were collected and analyzed using flow cytometry. The experiments were repeated for three times, and the proportion of cells in were shown in column graphs. **j-l.** Vero cells were synchronized to different stages and subsequently infected with TBEV, the mRNA level of TBEV envelope (E) gene was analyzed at 48 hpi by probe qPCR (**j**). The protein level of TBEV NS1 was analyzed at 48 hpi by immunoblot (**k, l**).

To elucidate the dynamic of TBEV-induced cell cycle arrest, we infected Vero cells with TBEV and monitored the cell cycle profile at 12, 18, 24, 36, and 48 hours post-infection (hpi). Our findings unveiled TBEV-triggered G0/G1 cell accumulation as early as 12 hpi, persisting throughout the 48 hour-infection period (Figure 4c). Furthermore, we noted a reduction in the percentage of cells in the S phase at 12 and 18 hpi, along with a decline in the proportion of cells in the G2/M phase at 48 hpi (Figure 4c), indicating that TBEV may initially impede S phase entry and later facilitate G2/M phase exit, ultimately leading to G0/G1 phase arrest.

Considering that cells in different phases of the cell cycle may impact virus-induced cell cycle arrest, we further investigated the effect of TBEV on synchronized cells. Cells were synchronized in the G0/G1, S and G2/M phases through serum starvation, thymidine and nocodazole treatments for 20 h (Figure S8e-g). The synchronized cells were then infected with TBEV, and their cell cycle progression was monitored (Figure S8g). In the case of serum-starved cells, the mock infection group displayed 58.48% of cells in the G0/G1 phase, a percentage that rose to 64.62% following TBEV infection (Figure 4d, e). Similar trends were observed in thymidine- and nocodazole-treated cells upon TBEV infection (Figure 4f-i), highlighting that TBEV can impact cell cycle progression, causing G0/G1 phase accumulation even in synchronized cells.

Given the critical role of cell cycle progression by viruses in promoting their replication, we sought to determine if G0/G1 phase synchronization enhances TBEV replication. Cells synchronized in G0/G1, S, and G2/M phases were infected with TBEV. At 2 hpi, serum starvation significantly increased TBEV RNA in infected cells, suggesting enhanced viral attachment or entry (Figure S8h). By 48 hpi, serum starvation resulted in elevated levels of TBEV RNA in both the supernatant and virus-infected cells (Figure 4j), whereas treatment with nocodazole or thymidine led to a slight reduction in viral RNA in virus-infected cells. Immunoblot and immunofluorescence analysis further showed that serum starvation significantly elevated the expression of TBEV NS1 protein, while thymidine and nocodazole treatment decreased NS1 protein expression, aligning with the trends observed in RNA content (Figure 4k, l, S8i), suggesting that TBEV exploits the G0/G1 cell cycle arrest to enhance its replication.

### TBEV NS5 induces G0/G1 cell cycle arrest via interacting with P300

To identify the viral protein responsible for TBEV-induced cell cycle arrest, we systematically evaluated the effects of individual TBEV proteins on cell cycle progression. Compared to the empty vector (EV) control, transfection with NS3, NS5, and prM significantly induced G0/G1 phase arrest. Notably, cells overexpressing NS5 displayed the most pronounced elevation in the G0/G1 phase cell population (Figure 5a, S9a). To validate NS5’s role in cell cycle regulation, we transfected cells with escalating doses of NS5 expression plasmids and analyzed the cell cycle profile. Dose-dependent enhancement of G0/G1 arrest was observed, with G0/G1 populations rose from 66.78% in the EV group to 70.71% and 75.52% in the 0.5 and 1 μg NS5 overexpression groups, respectively (Figure 5b, S9b).

**Figure 5.**
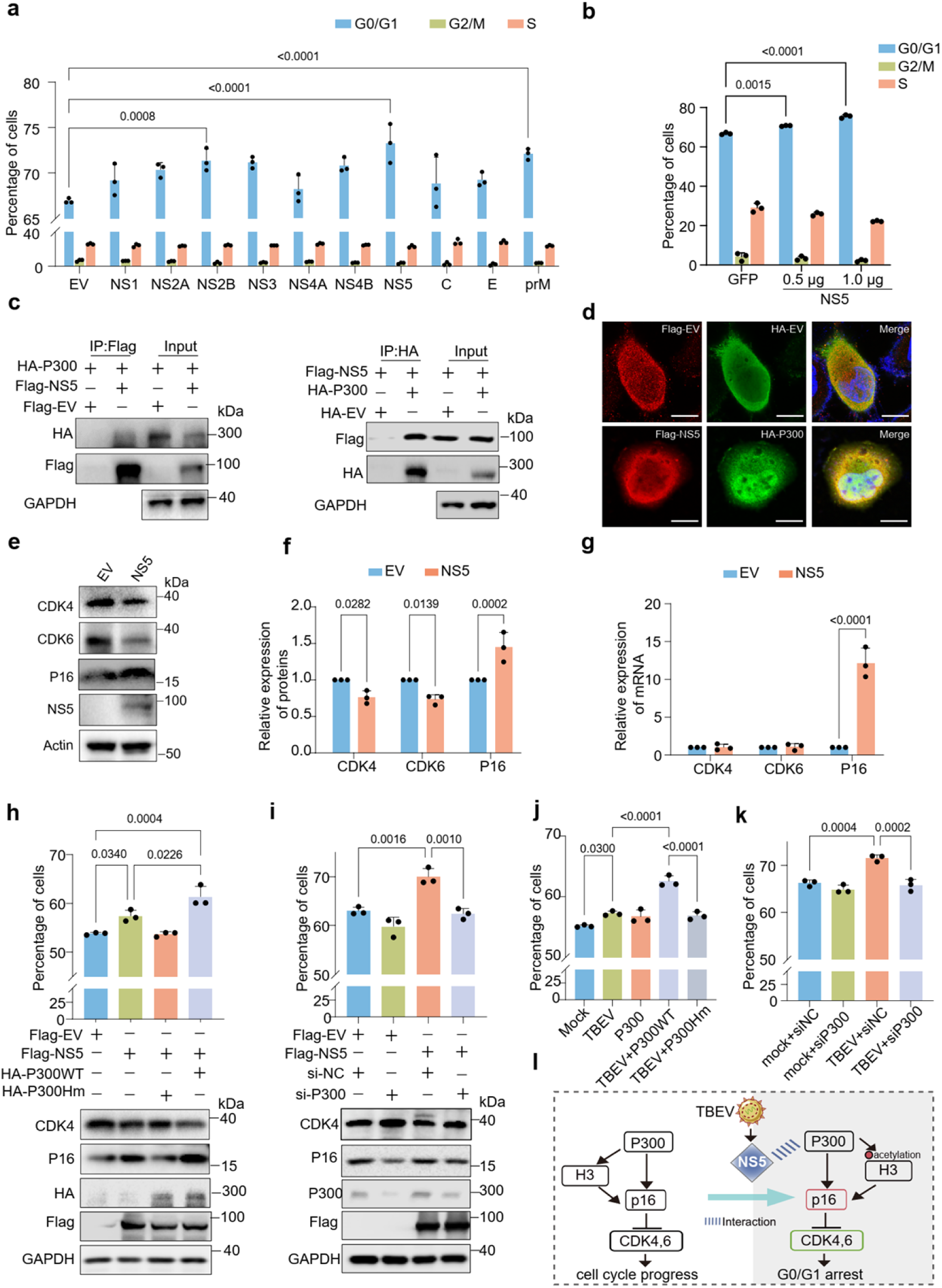
TBEV NS5 interacts with P300 to induce G0/G1 cell cycle arrest. **a.** Cells were transfected with the plasmids of Flag-tagged empty vector (EV) and ten TBEV viral proteins, the cell distribution was analyzed by flow cytometry. **b.** The Flag-EV, 0.5 and 1.0 μg Flag-NS5 were transfected into A549 cells, the cell distribution was analyzed by flow cytometry. The experiment was repeated for three times, and the percentage of cells were shown in column graph. **c.** Cells were transfected with Flag-EV or Flag NS5 together with HA-P300, the interaction between NS5 and P300 was analyzed by co-immunoprecipitation analysis. **d.** The colocalization of P300 (green) and NS5 (red) were examined by immunofluorescence analysis. Scale bar: 10 μm. **e-g**. Cells transfected with NS5 were collected after 48 h, the expression of CDK4, CDK6 and P16 was analyzed by immunoblot (**e**) and qPCR analysis (**f**). The relative expression of CDK4, CDK6 and P16 was shown in column graph (**g**). **h.** A549 cells transfected with HA-EV, HA P300 and P300 Hm were further transfected with NS5, the distribution of cells was analyzed by flow cytometry. The experiments were repeated for three times, and the proportions of cells were shown in column graph, the expression of indicated proteins was assessed by immunoblot assay. **i.** Cells transfected with P300 siRNA and NC siRNA (siNC) were further transfected with NS5, the distribution of cells was analyzed by flow cytometry, and the proportions of cells and the relative expression of P300 mRNA were shown in column graph. **j.** Cells transfected with HA-EV, HA P300 and P300 Hm were mock-infected or infected TBEV, the distribution of cells was analyzed by flow cytometry. **k.** Cells transfected with P300 siRNA and NC siRNA were mock-infected or infected TBEV, the distribution of cells was analyzed by flow cytometry. **l.** Outline of G0/G1 cell cycle arrest regulated by TBEV NS5.

To elucidate the mechanism underlying NS5-induced cell cycle arrest, we investigated P300, a histone acetyltransferase identified among NS5’s interaction partners and a known transcriptional activator of *P16*, a tumor suppressor that triggers G0/G1 arrest by inhibiting CDK4/6 and blocking G1 to S phase progression (*31*). We hypothesized that NS5 might engage P300 to augment P16 transcription, thereby suppressing CDK4/6 and inducing G0/G1 cell cycle arrest (Figure S7c). Co-immunoprecipitation analysis confirmed the interaction between NS5 and P300 (Figure 5c), further supported by their colocalization in immunofluorescence assays (Figure 5d). Overexpression of NS5 significantly increased P16 protein and mRNA expression while reducing CDK4/6 expression (Figure 5e-g). To elucidate P300’s functional role, we overexpressed wild-type P300 (P300WT) or a catalytically inactive mutant (P300Hm) lacking acetyltransferase activity. Intriguingly, P300WT amplified NS5-driven G0/G1 arrest, whereas P300Hm abolished this effect (Figure 5h, S9c). Similarly, P300WT co-expression synergized with NS5 to elevate P16 and suppress CDK4, while P300Hm attenuated these changes (Figure 5h). Conversely, siRNA-mediated P300 knockdown reversed NS5-induced P16 upregulation and CDK4 downregulation, restoring normal cell cycle progression (Figure 5i). Together, these results establish P300 as a critical mediator of NS5-induced G0/G1 arrest, dependent on its acetyltransferase activity.

To further validate the involvement of P300 in TBEV-induced cell cycle arrest, we overexpressed P300WT or P300Hm in TBEV-infected cells. Interestingly, P300WT exacerbated TBEV-triggered G0/G1 arrest, while P300Hm abrogated this phenotype (Figure 5j). Conversely, P300 knockdown markedly attenuated TBEV-induced cell cycle arrest (Figure 5k). Given that P300-mediated transcriptional activation of *P16* is associated with histone H3 acetylation, we assessed this modification. NS5 overexpression markedly increased H3 acetylation (Figure S9d), as did TBEV infection itself (Figure S9e). Taken together, these data demonstrate that TBEV NS5 hijacks P300’s acetyltransferase activity to transcriptionally upregulate *P16*, thereby suppressing CDK4/6 and enforcing G0/G1 arrest (Figure 5l).

### TBEV infection decreases CDK4 and upregulate P16 expression

We further validated the expression of CDK4, CDK6, P16 and cell cycle regulators following TBEV infection (Figure 6a). Consistent NS5 protein, TBEV infection induced a notable decrease of CDK4 and CDK6 at 36 and 48 hpi, while P16 exhibited increased expression at 36 hpi. Furthermore, the expression of CDK2, crucial for S phase progression, exhibited reductions at 12 and 24 hpi, aligning with the diminished percentage of cells in the S phase at 12 and 18 h upon TBEV infection (Figure 6a). Similarly, cyclin B1 and CDK1, pivotal for G2/M phase progression, displayed reduced expression at 36 and 48 hpi, mirroring the decline in the number of G2/M cells during the late infection phase (Figure 6a).

**Figure 6.**
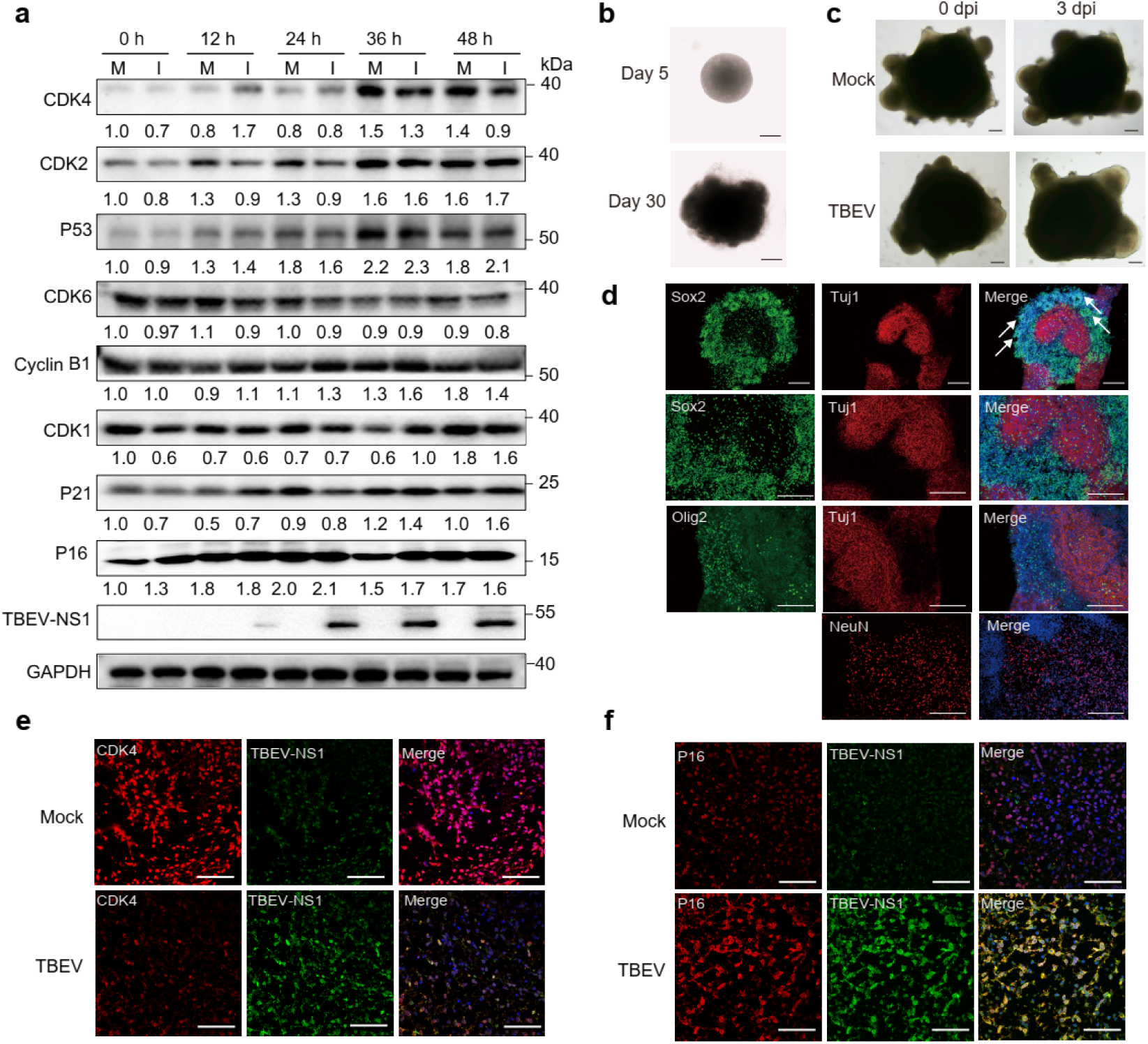
TBEV infection regulates the expression of CDK4 and P16 in brain organiods. **a.** Cells were either mock-infected or infected with TBEV, and then collected at 12, 24, 36 and 48 hpi, the expression levels of cell cycle regulators were detected by immunoblot using indicated antibodies. M: mock, I: infection. **b.** Representative images of brain organoid generation from iPSCs. Scale bar, 500 µm. **c.** Microscopic images of brain organoids, either mock-infected or infected with TBEV at 0- and 3-days post-infection (dpi). Scale bar, 200 µm. **d.** Microscopic images depicting 42-day-old iPSC-derived organoids, featuring TUJ1 (neurons), SOX2 (progenitors), Olig2 (oligodendrocytes), and NeuN (neuronal nuclei) markers. Scale bar, 500 µm. **e, f**. Brain organoids were either mock-infected or infected with TBEV at the MOI of 2.0, the expression of CDK4 (**e**) and P16 (**f**) in brain organoids were examined using immunofluorescence analysis. CDK4 and P16 were visualized in red, TBEV NS1was visualized in green, and the nuclei were stained in blue. Scale bar: 50 μm.

Brain organoids derived from induced pluripotent stem cells offer an exceptional platform for investigating neurotropic viruses. To delve deeper into the impact of TBEV on the expression of cell cycle regulators, we utilized TBEV-infected brain organoids (Figures 6b, c). Analysis of the brain organoids confirmed the presence of neural progenitor (SOX2), oligodendrocyte (Oligo2), immature neurons (TUJ1), and mature neuronal (NeuN) markers (Figure 6d). The detection of TBEV NS1 in the infected organoids validated successful virus infection (Figure 6e, f). Notably, compared to the mock infection group, TBEV infection led to a significant decrease in CDK4 expression and an increase in P16 expression (Figure 6e, f). These findings provide compelling evidence that TBEV infection disrupts the expression of CDK4 and P16, highlighting the dysregulation of cell cycle regulators following TBEV infection.

### Inhibitors of P300 and CDK4 agonist suppress TBEV replication

Given the pivotal role of the NS5-P300 interaction in cell cycle regulation, and the observed inhibitory effect of P300 knockdown on TBEV-induced G0/G1 arrest, we sought to investigate whether inhibitors of P300, C646 and CPI-637, could impede TBEV replication. Our results revealed that C646 exhibited more than 50% inhibition of TBEV replication even at the concentration as low as 10 nM in A549 cells, and displayed an IC_50_ of 1.46 μM in Vero cells (Figure 7a, S10a). Notably, CPI-637 demonstrated inhibitory effects on TBEV replication in both Vero and A549 cells without inducing significant cellular toxicity (Figure 7b, S10b). Subsequent immunoblot and immunofluorescence analyses demonstrated a substantial reduction in TBEV NS1 protein levels upon treatment with C646 and CPI-637 (Figure 7c-e), suggesting that these inhibitors have the potential to impede both TBEV replication and release. These findings strongly support the inhibitory effect of P300 inhibitors on TBEV replication.

**Figure 7.**
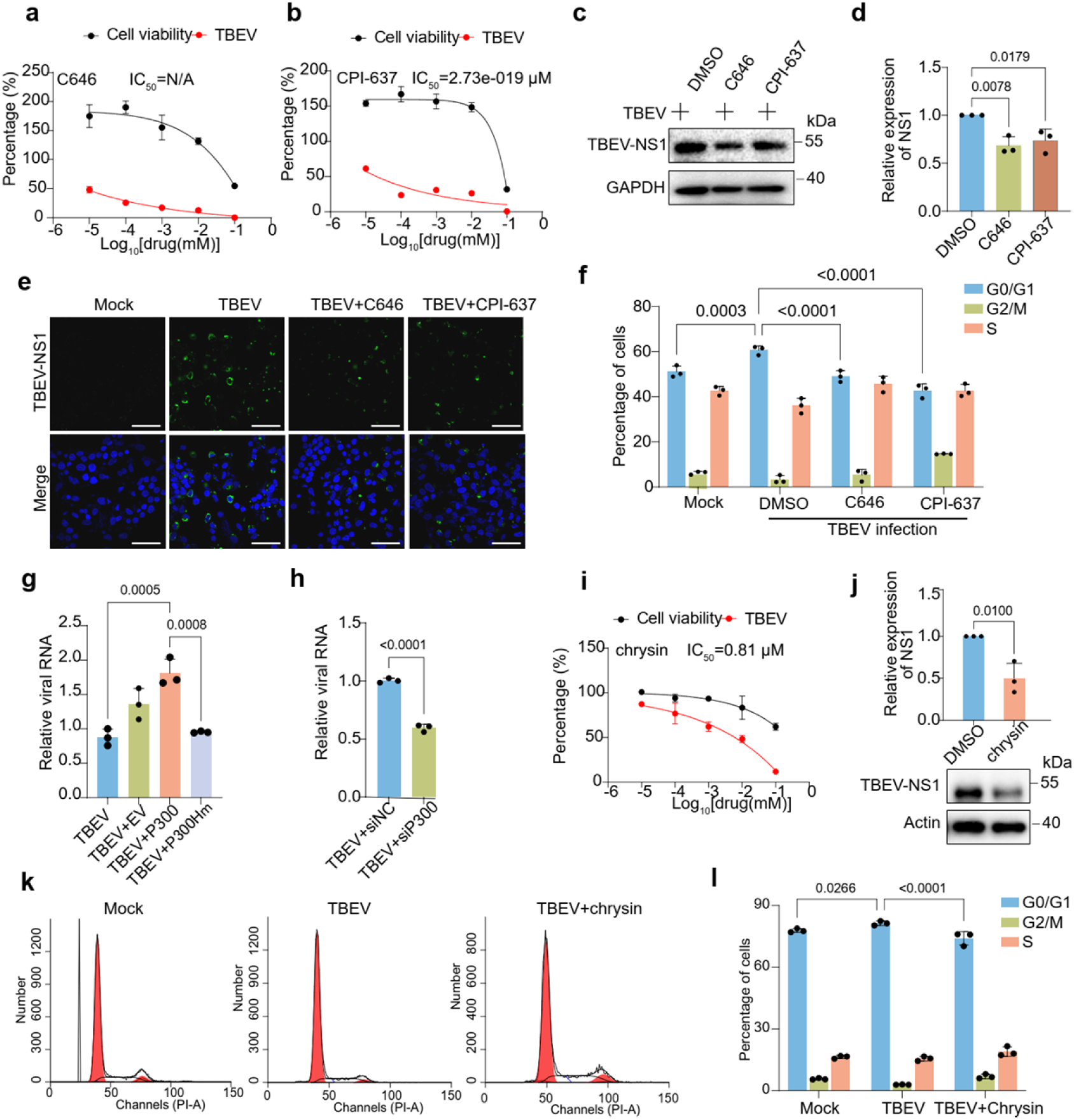
P300 inhibitors and CDK4 agonist restrict viral replication by interfering with TBEV-induced cell cycle arrest. **a, b**. A549 cells were either mock-infected or infected with TBEV, the cells were then treated with indicated concentration of C646 (**a**) and CPI-637 (**b**), the supernatants were collected at 48 hpi, and the mRNA level of TBEV E gene was analyzed by probe qPCR. **c-f**. Cells mock-infected or infected with TBEV at the MOI of 10 were treated with 10 μM C646 and CPI-637, the expression of TBEV NS1 protein was detected by immunoblot (**c, d**) and immunofluorescence analysis (**e**), and the cell cycle distribution was analyzed by flow cytometry (**f**). Scale bar: 50 μm. **g.** Cells transfected with HA-EV, HA P300 and P300Hm were infected TBEV, the mRNA level of TBEV E gene in the supernatants was analyzed by probe qPCR. **h.** Cells transfected with P300 siRNA and NC siRNA were infected TBEV, the mRNA level of TBEV E gene in the supernatants was analyzed by probe qPCR. **i.** A549 cells were treated with the indicated concentrations of chrysin for 2 h, the cells were then infected with TBEV, the supernatants were collected at 48 hpi, and the mRNA level of TBEV E gene was analyzed by probe qPCR. **j-l.** Cells were treated with 10 μM chrysin for 2 h, the cells were then mock-infected or infected with TBEV at the MOI of 10, the expression of TBEV NS1 protein was detected by immunoblot (**j**), and the cell distribution was analyzed by flow cytometry (**k**). The percentage of cells (**l**) upon chrysin treatment was shown in column graphs. Scale bar: 50 μm.

We then examined the cellular distribution of TBEV-infected cells treated with C646 and CPI- 637, revealing that C646 and CPI-637 significantly reduced the accumulation G0/G1 cells induced by TBEV infection (Figure 7f). This suggests that the repression effect of P300 inhibitors on TBEV replication may be mediated by interference with virus-induced cell cycle progression. Additionally, we assessed the impact of P300 on TBEV replication, demonstrating that P300 overexpression enhanced TBEV replication, while the mutant P300Hm exhibited no significant effect on viral replication (Figure 7g). Conversely, knockdown of P300 inhibited TBEV replication (Figure 7h).

Considering the observed decrease in CDK4 expression upon TBEV infection, we also evaluated the effect of chrysin, a natural flavone that can activate CDK4, on TBEV replication. Our findings revealed that chrysin effectively inhibited TBEV replication, displaying an IC_50_ of 0.81 μM in Vero cells and 0.78 μM in A549 cells (Figure 7i, S10c). Immunoblot and immunofluorescence analyses demonstrated that chrysin treatment significantly reduced the protein level of TBEV NS1 (Figure 7j, S10d). Additionally, chrysin treatment significantly reduced the accumulation of cells in the G0/G1phase induced by TBEV (Figure 7k, l). Collectively, these findings suggest that inhibiting P300 and activating CDK4 can suppress TBEV replication by disrupting virus-induced cell cycle progression.

### Identification of drugs that target host factors

To disrupt the TBEV interactome, we focused on identifying ligands for human proteins that interact with viral proteins, prioritizing interactions with MiST scores greater than 0.5. By comprehensively analyzing drug-target interactions from NeDRexDB (*32*), which integrates data from sources such as MONDO, DrugBank, and other biomedical databases, we identified promising therapeutic candidates. These compounds were selected on the basis of their development status (approved drugs or experimental candidates), target selectivity, and commercial availability. Network analysis revealed interactions between druggable human proteins and eight viral proteins, highlighting the interconnected nature of various cellular processes, including autophagy, ubiquitin-mediated proteolysis, ribosome biogenesis, and N-glycan/glycosaminoglycan biosynthesis. These findings suggest the potential for multitarget therapeutic strategies.

Through systematic screening and analysis of the virus-host interaction network, we identified several promising drug targets and their corresponding inhibitors (Figure 8a). Notably, our analysis revealed multiple druggable targets across diverse cellular pathways, such as EP300 in the cell cycle pathway (targeted by C646 and CPI-637), which has been demonstrated to inhibit TBEV replication (Figure 7a-e). Together with BRD9 in chromatin remodeling (targeted by sunitinib), and RIOK1 (inhibited by sunitinib/fostamatinib) in ribosome biogenesis. Sunitinib exhibited strong antiviral activity against TBEV, with an IC_50_ of 0.17 μM (Figure 8b), while fostamatinib showed significant effect on virus replication (Figure S10e). Additional therapeutic targets included XIAP (modulated by dequalinium chloride), and CDK9 in the HIV-1 infection pathway (targeted by abemaciclib). The X-linked inhibitor of apoptosis protein (XIAP) is an E3 ubiquitin-protein ligase that modulates hepatitis B virus replication via NF-κB signaling (*33*). In our study, XIAP was found to bind prM, and its inhibition using dequalinium chloride significantly suppressed TBEV replication, with an IC_50_ of 0.27 μM (Figure 8c). However, abemaciclib had no significant effect on TBEV replication (Figure S10f). Furthermore, we also identified the lysine acetyltransferases KAT6A and KAT7 (targeted by WM-1119 and WM-3835, respectively) that could interact with TBEV NS5. Inhibition of KAT6A using WM-119 significantly reduced the replication of TBEV, with an IC50 of 1.01 μM (Figure 8d). In contrast, WM-3835 that targets KAT7 exhibited less potent antiviral activity, with an IC_50_ of 1.16 μM (Figure S10g). Importantly, the deacetylase SIRT1 has also emerged as a particularly targetable node, with several available modulators, including SRT2183, resveratrol, fisetin, nicotinamide/nicotinamide chloride, and SRT1720 HCl markedly inhibited TBEV replication (*20*). This suggests that NS5 may regulate the acetylation levels of host proteins to promote viral replication.

**Figure 8.**
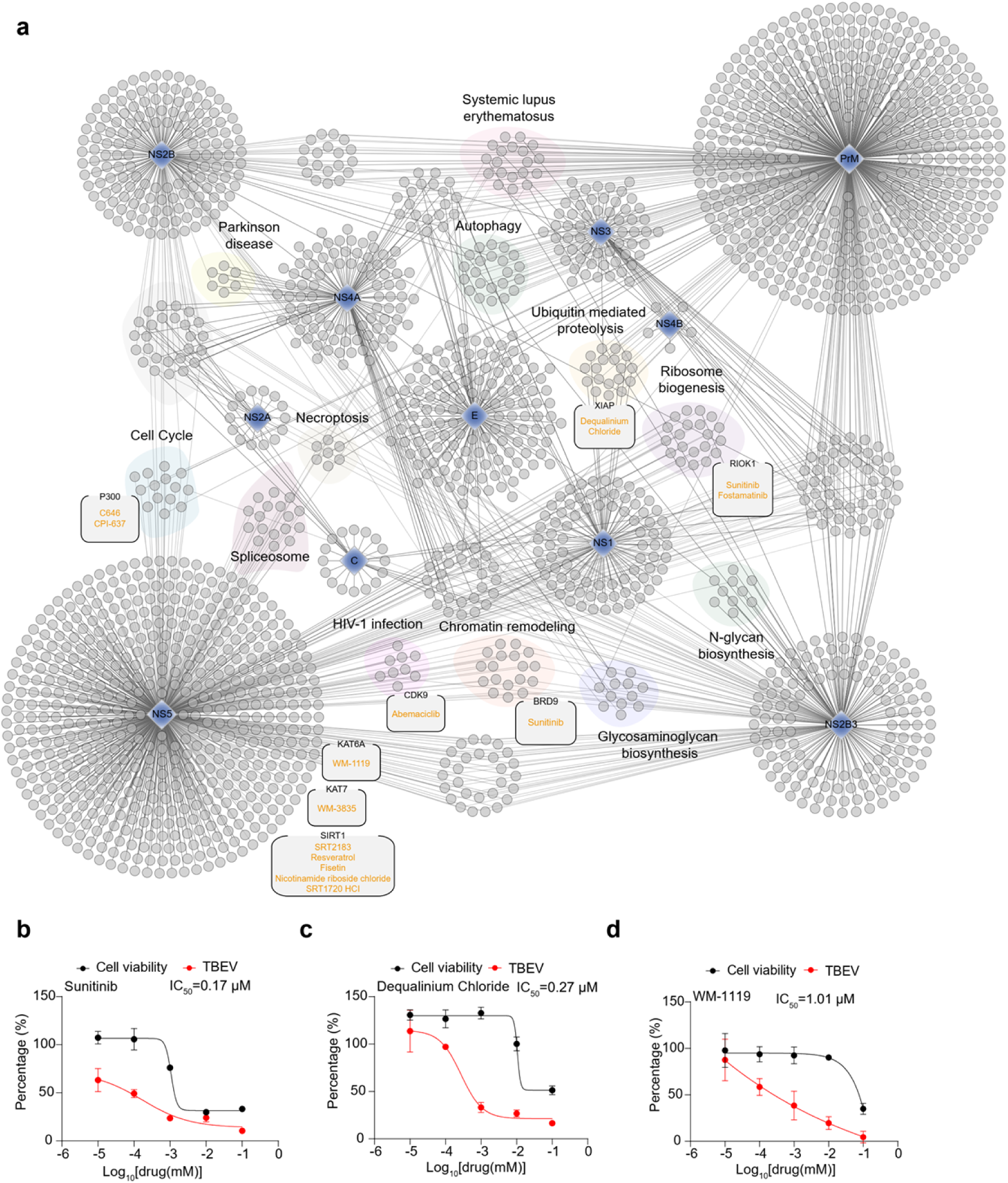
The TBEV interactome reveals pharmacological targets. **a.** Virus-protein interaction network and predicted drug targets. Potential drug-protein interactions were predicted by NeDREx(*32*) using the TrustRank algorithm, integrating data from various biomedical databases and experimentally validated targets. Nodes represent proteins and edges indicate their interactions. **b-d.** Cells were pretreated with indicated inhibitors targeting RIOK1/BRD9 (**b**), XIAP (**c**) and KAT6A (**d**) before TBEV infection for 48 h. The IC_50_ value was determined using probe qPCR. Cell viability was assessed using cell counting (CCK-8) assay in the absence of viral infection. The percentage of viral titer compared with DMSO treatment (red) and cell viability (black) depicted. Error bars represent the mean ± SEM of three biological replicates.

## Discussion

In this study, we employed a proteomics approach to systematically map TBEV-host protein interaction networks, uncovering viral reprogramming of critical cellular pathways, including innate immune signaling, spliceosome dynamics, DNA damage response, autophagy, and cell cycle regulation. Specially, we identified that NS5 protein directly interacts with histone acetyltransferase P300, mechanistically promoting P16 expression while downregulating CDK4, thereby triggering G0/G1 cell cycle arrest to promote viral replication. Our analysis further revealed potential antiviral targets, including P300-CDK4 axis, KAT6A, XIAP, and RIOK1/BRD9.

TBEV employs a sophisticated strategy to subvert the host’s interferon system. The viral NS5 protein disrupts interferon signaling through dual mechanisms: it directly binds prolidase, a protein essential for IFNAR1 surface expression (*34*), and interacts with Tyk2 kinase to block downstream signaling (*12*), a mechanism we have independently validated in this study. Notably, we identified a novel interaction between NS5 and SETD2, a methyltransferase that enhances interferon signaling by STAT1 methylation (*35*). Intriguingly, this parallels the immune evasion of SARS-CoV-2, whose nsp9 protein similarly inhibits STAT1-dependent signaling by targeting SETD2,(*36*) suggesting SETD2 may represent a conserved viral target for immune evasion. Ubiquitination of the K132 residue in TBEV NS4A was shown to be critical for its interaction with STAT1.(*14*) We further uncovered the novel NS4A-ESCIT interaction, shedding light on additional immune evasion strategies employed by NS4A. Consistent with prior reports (*15*), we confirmed that prM’s interaction with MAVS, which suppresses interferon production. Our study also newly links prM to TRIM4 and NS2B3 to UNC93B1, interactions that illuminate conserved viral strategies for evading host defenses. These findings collectively underscore the multifaceted mechanisms by which TBEV, and potentially other viruses, subvert antiviral signaling pathways.

Both DENV and ZIKV NS5 interact with host factors involved in RNA splicing (*19*). For example, DENV NS5 disrupts the cellular spliceosome by binding CD2BP2 and DDX23, core components of the U5 snRNP particle, thereby altering host RNA splicing to enhance viral replication (*37*). In our study, TBEV NS5 was found to interact with spliceosome-related proteins, such as CACTIN, ACIN1, RBM2, DHX8, and CDC40. Notably, ACIN1 and CDC40 were previously identified as ZIKV NS5 interactors (*19*), suggesting that conserved mechanisms among flaviviruses for co-opting mRNA splicing machinery.

Our findings uncover a multifaceted interplay between TBEV infection and host autophagy. Viral prM suppresses AKT-mTOR signaling via direct binding to AKT1, activating autophagy initiation (*20*). Additionally, prM and the NS2B3 protease regulate autolysosome maturation through interactions with VPS11 and STX17, respectively. Novel interactions, including prM- SPAG5, prM-Rab33B, and NS2B3-TSC2, further expand the roles of these viral proteins in modulating autophagic flux. The TBEV-host interactome also implicates NS1 and NS4B in autophagosome biogenesis. NS1 binds STK3 (a kinase regulating autophagy), while NS4B interacts with ATG7, a core autophagy protein critical for replication of hepatitis C virus (HCV) (*38*), influenza A virus (IAV) (*39*), and TBEV (*20*). The newly identified NS4B-ATG7 interaction may elucidate how TBEV exploits autophagy machinery, though the precise mechanism warrants further investigation.

Viral manipulation of the host cell cycle is a common pathogenic strategy (*40*). Enterovirus D68 induces G0/G1 arrest (*41*), while the Dabie bandavirus triggers G2/M arrest (*40*). Here, we demonstrate that that TBEV infection induces G0/G1 phase arrest, significantly enhancing viral replication. Conversely, nocodazole-mediated synchronization of cells at G2/M phase suppressed TBEV replication, aligning with prior studies (*42*). Flaviviruses broadly modulate cell cycle progression to enhance pathogenicity (*43, 44*). For instance, ZIKV infection induces S phase arrest by activating DNA damage response through ATM/chk2 pathway (*45*). JEV suppresses GSK3β expression, leading to prolonged cyclin D1 expression and subsequent G1 phase cell cycle arrest (*46*). Similarly, DENV infection triggers G1 phase arrest through the interaction of RNA hairpin in 3’ UTR with DDX6 (*45*). Consistent with JEV and DENV, TBEV infection also induces G0/G1 cell cycle arrest, characterized by decreased levels of CDK4 and CDK6, along with elevated P16 and P21 expression. Notably, we discovered that the NS5 protein, which encodes RdRp, plays a significant role in inducing G0/G1 phase arrest, aligning with the pivotal functions of RdRp proteins in modulating host cell cycle. For instance, the RdRp protein of HCV binds to CDK2 interacting protein (CINP), thereby inhibiting its nuclear localization and impeding S phase progression (*47*). Furthermore, RdRp proteins of SARS-CoV-1, SARA-CoV-2 and IBV interact with the P50 subunit of DNA polymerase δ to regulate host cell cycle progression (*48*). In the case of TBEV, the NS5 protein induces cell cycle arrest depending on the acetyltransferase activity of P300, indicating that the inhibition P300’s acetyltransferase activity could serve as a potential antiviral strategy. Additionally, other proteins of TBEV, including NS3 and prM, were also found to elevate the proportion of G0/G1 cells, warranting further investigation to comprehensively elucidate the mechanisms underlying TBEV-induced cell cycle arrest.

Drugs targeting virus directly can rapidly induce antiviral resistance. In contrast, host-directed antivirals, which rely on the pathogenic mechanisms of the virus, offer the dual benefits of a broad-spectrum efficacy and a lower risk of resistance (*49*). Based on TBEV-virus interactome, we revealed potential antiviral targets, including P300-CDK4 axis, KAT6A, XIAP, and RIOK1/BRD9. Chrysin, a dietary phytochemical, exerts potent inhibition on enterovirus 71 replication by suppressing viral 3Cpro activity (*50*). Additionally, chrysin has been reported to inhibit influenza virus replication of by repressing virus-induced cell cycle arrest and apoptosis (*51*). The P300 inhibitor, C646, has demonstrated significant inhibitory effects on the replication of herpes simplex virus-1, respiratory syncytial virus, and various strains of influenza virus (*52–54*). Furthermore, CPI-637 functions as a dual-target inhibitor to antagonize HIV-1 latency (*55*). Sunitinib, the inhibitor of RIOK1/BRD9, showed antiviral effect on ZIKV virus (*56*). The exploration of compounds targeting these specific proteins may present promising avenues for the development of antiviral therapies against TBEV and other related infections.

In summary, our study systematically maps the TBEV-host interactome, revealing viral subversion of critical pathways, including innate immune signaling, spliceosome dynamics, DNA damage response, autophagy, and cell cycle regulation. Mechanistic insights into NS5-mediated G0/G1 arrest and novel interactions advance understanding of TBEV pathogenesis. Furthermore, the identification of host-directed antiviral candidates underscores the translational potential of targeting virus-host interfaces. These findings provide a roadmap for developing therapies against TBEV and related flaviviruses.

## Materials and methods

### Cells and viruses

African green monkey kidney epithelial cell (Vero), human lung cancer alveolar basal epithelial cell (A549), human brain glioma cell line (T98G), and human embryonic kidney cells (HEK293T) were cultured in DMEM (HyClone, Logan, USA) supplemented with 10% FBS (BBI, Shanghai, China) and 1% penicillin-streptomycin (Pen/Strep). All cells were incubated at 37°C in a humidified atmosphere with 5% CO_2_.

The TBEV virus, classified as the far-eastern type, was isolated in northeastern China.(*57*) The virus was propagated in Vero cells, and the titer of virus was assessed using tissue culture infective dose (TCID_50_) method.

The target cells were rinsed with serum-free medium, followed by the addition of the virus in the serum-free medium at the designated multiplicity of infection (MOI). After a 2-hour incubation period, the medium was replaced with DMEM supplemented with 10% FBS and 1% Pen/Strep. All experiments involving TBEV infection were conducted in a Biosafety Level 3 (BSL-3) laboratory.

### Plasmids, siRNA, antibodies and drugs

The genes encoding both the structural and non-structural proteins of TBEV (Genbank access number MN615726.1) were cloned into a Flag-VR1012 expression vector from Sangon Biotech (Shanghai, China). HA-p300 and HA-P300 Hm (the mutant form of P300 lacking histone acetyltransferase activity) plasmids were purchased from MiaoLing Bio (Wuhan, China). For small interfering RNA (siRNA) targeting P300, 5’-GGACUACCCUAUCAAGUAATT-3’, was synthesized at Sangon Biotech (Shanghai, China). Antibodies against HA, Flag, GAPDH, Actin, CDK1, CDK2, CDK4, CDK6, p16, p21, p53, Cyclin B1, TBEV-NS1, as well as CoraLite488- conjugated goat anti-mouse IgG antibody were obtained from Proteintech (Wuhan, China). Thymine, nocodazole, chrysin, the antiviral drugs were sourced from Selleck, USA.

### Affinity purification of viral proteins

To perform affinity purification of viral protein, 10 million HEK239T cells were transfected with 25 μg individual viral plasmids using PEI transfection reagents (Yeason, China) according to the manufacturer instruction. For each bait, three to four independent biological replicates were prepared. Cells were collected at approximately 48 hpi and lysed using immunoprecipitation (IP) buffer (50 mM Tris (pH 7.5), 1 mM EGTA, 1 mM EDTA, 1% Triton X-100, 150 mM NaCl), supplemented with 100 µM phenylmethylsulphonyl fluoride (PMSF) and TM protease inhibitors (Selleck, Houston, USA). The samples were incubated at 4°C for 30 min, and the debris was pelleted by centrifuge at 4°C,12,000 rpm for 15 min. Supernatants were collected, and viral and its interacting host proteins were purified using Strep-Tactin^®^ XT Purification column (IBA Lifesciences, Germany). Briefly, after column equilibration, the supernatants were added to the column and washed five times with 1 ml wash buffer. Proteins were eluted using 3 ml elution buffer containing 50 μM biotin, and the elutes were then concentrated using Amicon Ultra-4 (Millipore, USA). All wash and purification steps were conducted at 4℃. The expression of Strep-tagged proteins was analyzed by immunoblotting and Coomassie brilliant blue stain, respectively.

### Mass spectrometry analysis

Enriched proteins underwent reduction and alkylation using 10 mM dithiothreitol (DTT) and 55 mM iodoacetamide (IAM), respectively. The proteins were then digested and analyzed using the nanoflow reversed-phase liquid chromatography System (EASY-nLC 1200, Thermo Scientific) coupled to the Orbitrap Exploris 240 Mass Spectrometer (Thermo Scientific). The reverse-phase peptide separation was conducted using a trap column (75 μm ID × 2 cm) filled with 3 μm 100 Å PepMap C18 medium (Thermo Scientific), followed by a reverse-phase column (25 cm long × 75 µm ID) packed with 2 µm 100 Å PepMap C18 medium (Thermo Scientific).

The mobile phase consisted of solvent A (0.1% formic acid (FA) in water) and solvent B (0.1% FA in 80% acetonitrile (ACN)). Peptides were separated by loading them into the trap column at a flow rate of 5 µl/min and subsequently eluted from the analytical column at a flow rate of 300 nl/min. The liquid chromatography (LC) gradient was applied as a linear gradient from 8% to 27% of solvent B over 72 min, followed by a transition from 27% to 36% over the next 18 min. Subsequently, the gradient was rapidly increased to 90% within 2 min and held at that level for 10 min. The mass spectrometer was operated in positive ion mode with standard data-dependent acquisition, and the Advanced Peak Detection function was enabled for the top 20 ions. Precursor ion fragmentation was achieved using a normalized collision energy of 30% via High Collision Dissociation (HCD). The Orbitrap mass analyzer was set to a resolution of 60,000 for MS1 and 15,000 for MS2. Full scan MS1 spectra were collected in the mass range of 350-1,400 m/z, with an isolation window of 1.2 m/z and a fixed first mass of 100 m/z for MS2. The spray voltage was set at 2.2 kV, and the Automatic Gain Control (AGC) target was set at 300% for MS1 and the standard setting for MS2, respectively. The maximum ion injection time was set to 50 ms for MS1 and 110 ms for MS2.

### Analysis of protein-protein interaction

For the processing of affinity purification and mass spectrometry (AP-MS) data, MaxQuant software (version 1.6.17.0) was employed using default settings, including the label-free quantification (LFQ) method for analyzing protein intensities. The protein databases used comprised human sequences (20,407 proteins, downloaded on May 25, 2021), UniProt-reviewed Mycoplasma sequences (3,040 proteins, downloaded on May 25, 2021), and TBEV proteins. LFQ intensity served as the primary metric for protein quantification, enabling the comparison between experimental and control groups. To ensure the reliability of the results, protein identification and peptide-spectrum match (PSM) validation were rigorously maintained at a false discovery rate (FDR) of 1%.

Each TBEV bait MS dataset underwent scrutiny using the Mass spectrometric Interaction STatistics (MiST) algorithm.(*58*) The MiST algorithm assigned scores to each identified protein, predicting their likelihood of being TBEV interacting partners. Proteins with TBEV bait-predator MiST scores of >0.5 were identified as specific interactors with TBEV.

### Comparative analysis of TBEV and flaviviruses interactomes

To further analyze the TBEV-host interactome, we conducted a comparative analysis by incorporating three previously reported interactomes of other flaviviruses, namely DENV, ZIKV, and WNV. This assessment was conducted using UpSet R package,(*18, 19*) which specializes in evaluating the overlap of virus-host protein-protein interactions. The UpSet R package enabled the visual representation of intersections and unique components among multiple datasets. this approach facilitated a clearer understanding of the similarities and differences between various interactomes.

### Enrichment and Network Analysis

For the AP-MS data, enrichment analysis was conducted using the clusterProfiler R package to perform both Kyoto Encyclopedia of Genes and Genomes (KEGG) and Gene Ontology (GO) analyses, with an adjusted P-value threshold of < 0.05 for downstream analysis. Network analysis was carried out using the STRING database (https://string-db.org/), with visualizations created in Cytoscape (version 3.10.3). The regulated virus-host interactions, as well as host protein modifications including phosphorylation and acetylation, were illustrated using the pathview R package.

### Cell cycle analysis by flow cytometry

Following collection, the cells were fixed with 70% ethanol at 4 °C for 20-24 h. Subsequently, the fixed cells were washed with PBS and stained with PI/RNase Staining Buffer Solution (BD Biosciences, California, USA) for 15 min. Cell clumps were removed, and cell cycle distribution was assessed using a BD LSRFortessa flow cytometer. The resulting data were analyzed using ModFit software.

### Cell synchronization

To synchronize cells in the G0/G1 phase, serum deprivation was carried out for 20 h. For S phase synchronization, cells were treated with a final concentration of 0.85 mM Thymi in the growth medium for 20 h. To synchronize cells in the G2/M phase, 25 ng/ml of nocodazole was added to the growth medium for 20 h.

### Generation of brain organoids

Brain organoids were created using the brain organoid generation kit from Guidon Pharmaceutics (Beijing, China). In summary, induced pluripotent stem cells (iPSC) were dissociated into a single-cell suspension using accutase. The cells were then seeded into PrimeSurface low-adhesion 96-well plates. Once the embryoid bodies (EBs) reached a diameter of 500-600 μm with distinct boundaries, they were transferred to a low-adhesion 24-well plate. Following 4 days of static culture using medium without Vitamin A, the organoids were moved to a 37°C shaking incubator and cultured with medium containing Vitamin A until further use. This study received approval from the Medical Ethics Committee of the First Affiliated Hospital of Jilin University (2023–661).

### Coimmunoprecipitations and immunoblot assay

The cells were lysed in lysis buffer (50 mM Tris-HCl, pH 8.0, 150 mM NaCl, 1% NP-40) supplemented with a protease and phosphatase inhibitor cocktail. For co-immunoprecipitation, the lysates were incubated overnight with anti-flag M2 affinity gel (Sigma-Aldrich, St Louis, USA). Subsequently, the proteins were separated by SDS-PAGE and transferred onto PVDF membranes. Following blocking with PBST containing 5% BSA, the membrane was incubated overnight at 4°C with the primary antibody. The membrane was then thoroughly washed with PBST before the addition of a secondary antibody corresponding to the primary antibody. The protein bands on the membrane were visualized using a chemical Doc XRS molecular imager software (Bio-Rad, Philadelphia, PA, USA) with ECL luminescence solution.

### Detection of histone H3 total acetylation

The detection of histone H3 total acetylation was performed using the Histone H3 Total Acetylation Detection Fast Kit (Abcam, USA). The extracted total histones, along with the standard control, were added to the plates. Subsequently, detection antibodies were added to each well after washing, followed by the addition of the color developer solution. The absorbance was then measured on a microplate reader at 450 nm.

### Quantitative PCR (qPCR) and probe qPCR

Total RNA was isolated using the EasyPure® RNA Kit (TransGen, Beijing, China), followed by cDNA synthesis with the transScript first-strand cDNA synthesis super mix (TransGen, Beijing, China). Quantitative PCR (qPCR) was conducted using SYBR Green Master (Roche, Basel, Switzerland). The expression level of P16 was assessed using the specific primers: forward primer for P16 was 5’- CCGAATAGTTACGGTCGGAGG-3′, reverse primer: 5’- CACCAGCGTGTCCAGGAAG-3’. The forward primer for P300 was 5’- TGCGATCTGATGGATGGTCG-3’, the reverse primer was 5’- ACAGACAGTACAGTGCCAGC-3’. Data were normalized to the house keeping gene GAPDH.

Viral RNA was detected by probe-based qPCR. RNA was extracted using the TIANamp Virus RNA kit (Tiangen, Beijing, China) following the manufacturer’s protocol. Subsequently,viral RNA was reverse-transcribed to cDNA, and the copies of TBEV were quantified with specific primers: forward primer 5′-GGGCGGTTCTTGTTCTCC-3′, reverse primer 5′- ACACATCACCTCCTTGTCAGACT-3′, and the probe FAM- TGAGCCACCATCACCCAGACACA-BHQ1.

### Immunofluorescence analysis

The cells were initially fixed with 4% paraformaldehyde (PFA) for 30 min in the dark, followed by permeabilization with pre-cooled PBS containing 0.5% TritonX-100. Subsequent to fixation, the cells were washed with PBST and blocked with a 1% BSA blocking solution at 37°C for 1 h.The cells were then stained with primary antibody, followed by secondary antibody staining, and counterstained with DAPI (Yesen Biotechnology, Shanghai, China) for nuclei visualization. Finally, fluorescent images were acquired and analyzed using a confocal microscope (FV3000, Olympus).

### CCK8 detection of cell viability

The cells were seeded in 96-well plate and allowed to adherence for 24 h. Subsequently, drugs at varying concentrations were diluted in culture medium and used to treat the cells for 48 h. Each dilution was performed in triplicate. Following the incubation period, 10 μl of CCK8 solution was added to each well and incubated for 4 h in a cell culture incubator. The optical density (OD) value was measured at 450 nm using a microplate reader. Cell viability was quantitatively analyzed based on the measured OD values.

### Statistical analysis

Statistical analysis was performed using GraphPad Prism 9 software. The t-test was utilized for comparisons between two groups, while analysis of variance (ANOVA) was employed to assess statistical differences among multiple groups. Data were presented as mean ± standard deviation (SD), with statistical significance set at *P* < 0.05.

## Author contributions

Conceptualization: LS, NL, YZ, QL

Methodology: HW, YZ

Investigation: YHZ, HC, HX, TT, WW, MZ, NZ, ZS, YX, KZ, LC, GW

Visualization: HW, YZ

Supervision: YZ, QL

Writing—original draft: LS

Writing—review & editing: LS, QL

## Data and materials availability

All data are available in the main text or the supplementary materials. The mass spectrometry data are available in ProteomeXchange under the identifier PXD048439.

